# Theta-phase dependent neuronal coding during sequence learning in human single neurons

**DOI:** 10.1101/630095

**Authors:** Leila Reddy, Matthew W. Self, Benedikt Zoefel, Marlène Poncet, Jessy K. Possel, Judith C. Peters, Johannes C. Baayen, Sander Idema, Rufin VanRullen, Pieter R. Roelfsema

## Abstract

The ability to maintain a sequence of items in short-term memory (STM) is a fundamental cognitive function. In the rodent hippocampus, the representation of sequentially organized spatial locations is reflected by the phase of action potentials relative to the theta oscillation (phase precession). We investigated whether the timing of neuronal activity relative to the theta brain oscillation also reflects sequence order in the medial temporal lobe of humans. We used a task in which human subjects learned sequences of pictures and recorded single neuron and local field potential activity with implanted electrodes. We report that spikes for three consecutive items in the sequence (the preferred stimulus for each cell, as well as the stimuli immediately preceding and following it) were phase-locked at distinct phases of the theta oscillation. Consistent with phase precession, spikes were fired at progressively earlier phases as the sequence advanced. These findings generalize previous findings in the rodent hippocampus to the human temporal lobe and suggest that encoding stimulus information at distinct oscillatory phases may play a role in maintaining their sequential order in STM.

## Introduction

Learning to memorize and maintain a sequence of items in short-term memory (STM) is fundamental for successful behavior. At the cellular level, this process is linked to sustained firing activity of individual neurons during the delay period between two stimuli ^1-3^, as well as by anticipatory activity before the onset of an expected stimulus ^4^. Short-term memory processes are also linked to various brain rhythms ^5,6^, and there is converging evidence that neuronal firing activity during short-term memory can be timed (or phase-locked) to theta oscillations. In humans, the strength of theta phase locking is predictive of human memory strength ^7^ and navigational goals ^8^. In rodents, spiking activity of place cells ^9^ is locked to specific phases of the theta rhythm during spatial navigation. As a rat runs through a sequence of spatial positions, place cells that represent each of these locations fire at distinct phases of the underlying theta rhythm, and their spikes occur at increasingly early phases as the rat approaches the place fields. This process has been called phase-precession and has been proposed to play an important role in the learning of a sequence of spatial positions in the rodent brain ^10^ and it may, by extension, also play a role in the encoding and maintenance of any ordered list in STM ^11^.

Here we asked whether a theta phase-dependent coding scheme is also observed in the human brain during sequence learning. Because humans are more visual creatures than rodents, we assumed that learning a sequence of visual objects could be analogous to learning a sequence of spatial positions in rodents. We hypothesized that theta phase could play an important role in stimulus encoding, and that the phase at which a given cell fires would vary with stimulus identity and order. In other words, while subjects are involved in learning a stimulus sequence, each item in the sequence might be represented by neuronal activity that is locked to a different theta phase. Will medial temporal lobe (MTL) neurons in human subjects that navigate a “conceptual” space, defined by a sequence learning task, exhibit the same form of phase precession that has been observed in rodents during spatial navigation? Do MTL neurons also fire at increasingly early phases when the subject approaches the concept that best activates the cells?

## Results

To create an analogy with the navigation of spatial positions in rodents, we designed a conceptual space that consisted of a sequence of images. We defined a conceptual space in which images were displayed on the rim of a rotating “wheel” that moved in the clockwise direction. The wheel moved forward smoothly during the inter-stimulus interval (ISI; 0.5 seconds), during which period, a gray placeholder covered all the images. At the end of the ISI period, the wheel stopped for 1.5 seconds, and the placeholder at the topmost position of the wheel was replaced by the next image in the sequence (Figure 1A). We incentivized the subjects to learn the sequences by including probe trials (20% of trials). On probe trials we did not present the next stimulus, but showed two choice stimuli instead, and the observers had to indicate which stimulus was the next in the sequence. In our previous work with the same paradigm ^4^, we showed that human MTL neurons that initially responded to a particular (“preferred”) image on the wheel started firing in anticipation of this preferred image as a result of sequence learning, during the immediately preceding stimulus and the intervening ISI. This finding is reminiscent of rodent place cells that show anticipatory activity in sequentially ordered spatial environments ^12^. In the current study, we ask whether theta-phase coding mechanisms observed in rodent place cells also occur while humans navigate this conceptual space. In other words, are different stimuli in the sequence assigned a particular theta phase for firing, and is the order of theta phases similar to that observed in rodent phase precession?

**Figure 1:**
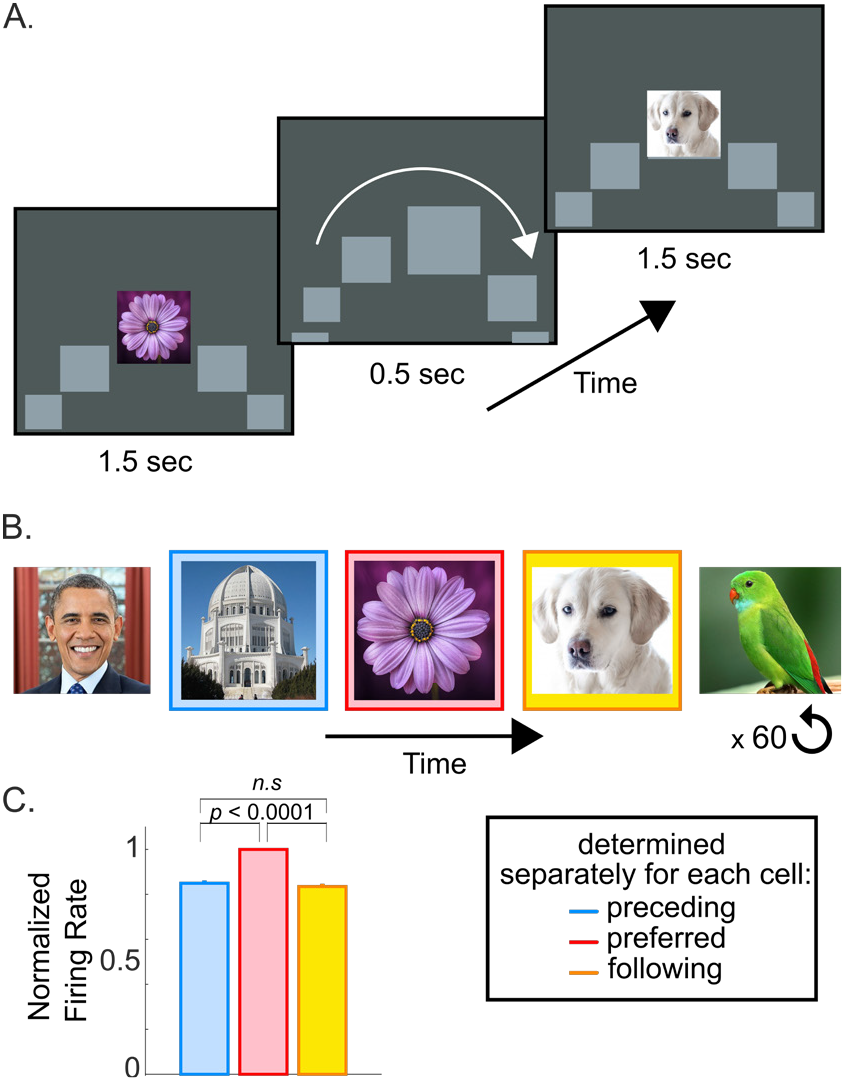
Experimental Design. A) In the sequence learning experiment, a sequence of 5-7 images was presented to the subjects in a fixed order. Each image was presented for 1.5s followed by an ISI of 0.5s. During the image presentation period (1.5s), an image was presented at the center of the screen, flanked by gray placeholders. During the ISI period, a placeholder replaced the central image, and all the placeholders moved in the clockwise direction. At the end of the ISI period, the central placeholder was replaced by the next image in the sequence. B) An example of a 5-image sequence. The sequence was repeated 60 times in each experimental session. C) For each neuron that we recorded from (*N* = 619) we determined the image that elicited the most spikes. This image was designated as the preferred image for that neuron. The images before and after the preferred image in the sequence were designated as the preceding and following images, respectively. The preferred image had the highest firing rate, which was significantly different from the neuronal response to the preceding and following images (*p*<0.00001, paired t-test performed on cross-validated data; see main text); the responses to the preceding and following images were not significantly different from each other (*p*=0.96 paired t-test). Error bars correspond to the standard error of the mean (*SEM*) across cells.

Nine human subjects learned the order of a fixed number of stimuli presented in a pre-defined sequence, while we recorded spiking and LFP activity from 619 neurons in the hippocampus, parahippocampal gyrus, and temporal cortex. Subjects rapidly learned the sequence order (>90% performance on test trials within 6 sequence presentations ^4^), and consequently all trials except the probe trials were included in the analyses. For each neuron we identified the stimulus that elicited the largest number of spikes (i.e., the “hot spot” in the sequence) and designated this stimulus as the preferred stimulus. The stimuli before and after the preferred stimulus were labeled as the preceding and following stimuli, respectively (Figure 1B, C; see below for results based on a cross-validated analysis). Here, we compared the theta-phase of firing for the preceding, preferred and following stimuli, and determined whether neuronal responses elicited by these stimuli are encoded at distinct theta phases, reflecting the order in which the sequence unfolds.

In accordance with previous studies ^7,8,13,14^, we observed prominent oscillatory activity in the LFP power and the spike-triggered power (STP) across all recording electrodes in the theta (4-8Hz) and beta (10-18Hz) bands (Figure 2 A, B). Neuronal spiking activity was significantly phase-locked (Raleigh test, *p*<0.00005) in both frequency bands (Figure 2C), with a preferred firing phase around the trough of the oscillation (pi radians).

**Figure 2:**
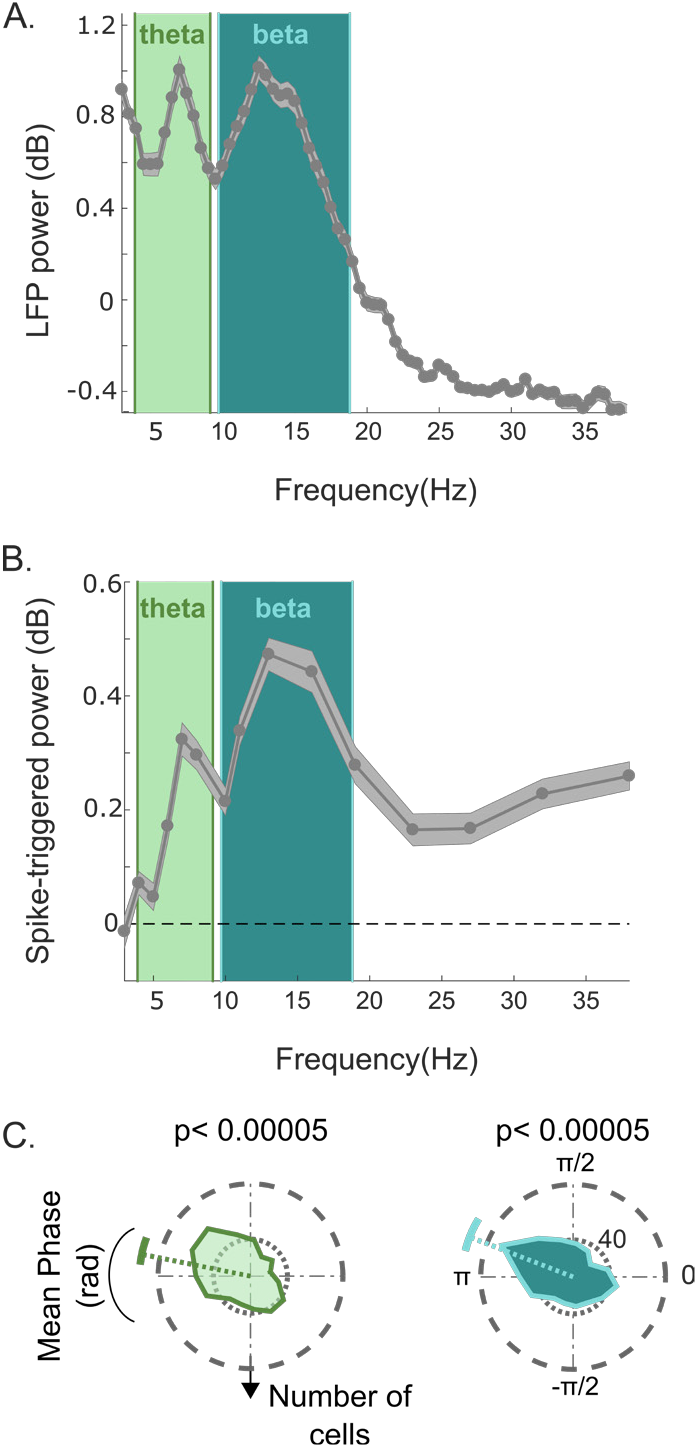
LFP power and theta and beta phase-locking of neurons. A) LFP power (expressed in decibel units relative to a 1/f fit, see Methods) reveals prominent activity in the theta (4-8Hz) and low-beta (10-18Hz) bands. B) Theta and beta oscillations in the power spectra of the spike triggered power (STP) during the sequence learning sessions (relative to a 1/f fit). Again, peaks are observed in the theta and beta bands. In A) and B), the solid line and shaded area correspond to the mean and *SEM* across all recording channels, respectively. C) Distribution of preferred firing phases across cells in the sequence learning sessions with respect to the theta and beta oscillations. The colored dotted and angular solid lines respectively correspond to the mean and standard deviation of the preferred firing phase across cells. The p-values are from a Rayleigh test.

To determine whether the neurons encoded the preceding, preferred, and following stimuli at different phases of the theta and beta cycle during sequence learning, we next compared the phase of firing for these stimulus types. The neurons fired at distinct phases for these stimuli in the theta band (Figure 3A, B). Phase coding did not occur in the beta band (Figure 3C, D) despite a strong beta band oscillatory component (Figure 2C). To determine significance, we applied a cross-validation procedure in which we based the neuron’s preferred stimulus on all trials except one and analyzed phase-coding in the left-out trial (see Methods and Figure S1). The pair-wise difference in phase preferences for the different stimulus types was significant across cells (F(1,1241) = 24.0; p<0.0001 for preferred vs. preceding; F(1,1241) = 29.4; p<0.0001 for preferred vs. following; and F(1,1241) = 105.18; p<0.0001 for preceding vs. following; Watson-William test Bonferroni-corrected for multiple comparisons).

**Figure 3:**
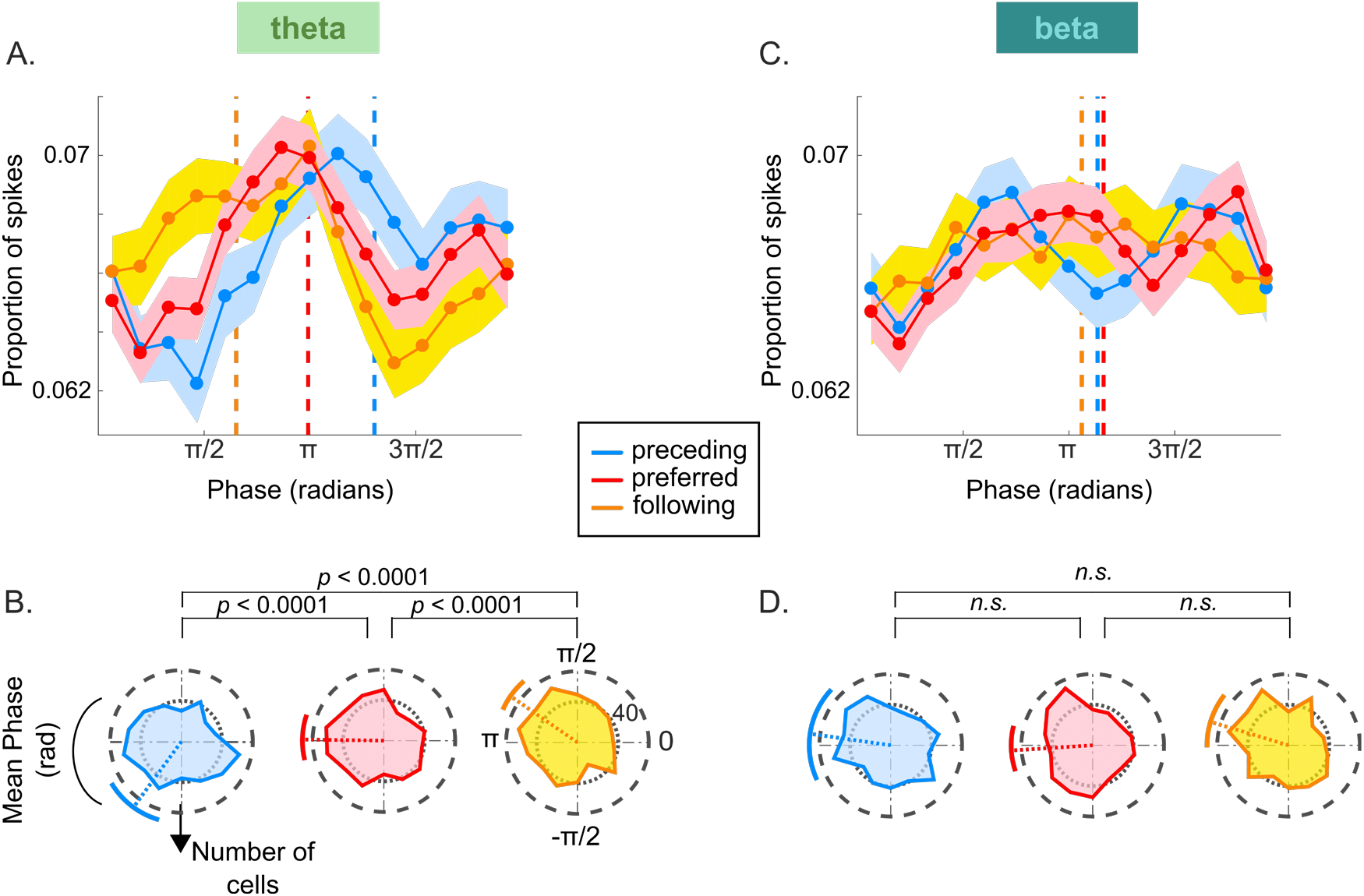
Phase encoding of successive stimuli. A, Distribution of spike phases relative to the theta band LFP for the preceding (blue), preferred (red) and following (yellow) stimuli across cells (*N* = 619). The dashed colored lines indicate the mean phase, the shaded areas correspond to the *SEM* across cells. The spike distributions have been smoothed for display purposes, but the mean phase (dashed lines) is calculated on unsmoothed data. B, Circular histogram of the firing phase across all cells (*N=619*), with colors as in panel (A). The dashed colored lines indicate the mean phase, and the colored angular lines correspond to the standard deviation across cells. The phase distributions were significantly different from the uniform distribution, indicating phase locking (*p* <0.05 for all stimulus types, Rayleigh test). The p-values above the rose plots are the results of a multi-sample Watson-William test for equal means, Bonferroni-corrected for multiple comparisons. C, D, Spike phase distribution and preferred phase histograms relative to the beta band oscillation for the three stimulus types. The neurons showed significant phase locking only for the preferred and following stimuli (p<0.05, Raleigh test). In contrast to the theta band, the cells did not fire at distinct phases for the different stimuli. The format of these panels is similar to panels A and B.

Importantly, similar to the phenomenon of phase precession in the rodent hippocampus, phases advanced to earlier values as the sequence progressed (mean angle ± standard deviation = 234°±23° (preceding), 178°±18° (preferred), and 118°±14° (following). The difference of ~58° between successive stimuli corresponds to a time difference of ~28ms with respect to a 6 Hz oscillation. Phase precession implies that the cycle of action potentials is slightly shorter than the theta band cycle in the LFP. It predicts a slightly higher frequency in spiking compared to the LFP theta oscillation ^15^ and this is indeed what we observed (Figure S2). To examine how the preferred phase evolved at a finer time scale as the sequence progressed from one stimulus to the next, we plotted the mean phase across cells as a function of time (Figure 4). We observed distinct phase-of-firing during the time periods in which the stimuli were presented, and a phase transition in between stimuli. A linear fit across cells through the phase values during stimulus presentation revealed a negative slope in the theta band (linear regression slope = -22.9 deg/s, r^2^ = 0.92, *p* <0.0005), but not in the beta band (slope = 1.2 °/s, r^2^ = 0.02, *p* = 0.4). The result was confirmed when we examined the phase slopes of individual cells, which were significantly negative in the theta band (t(613)=-4.8, p<0.00005), but not in the beta-band (t(613)=-0.2, p>0.5).

**Figure 4:**
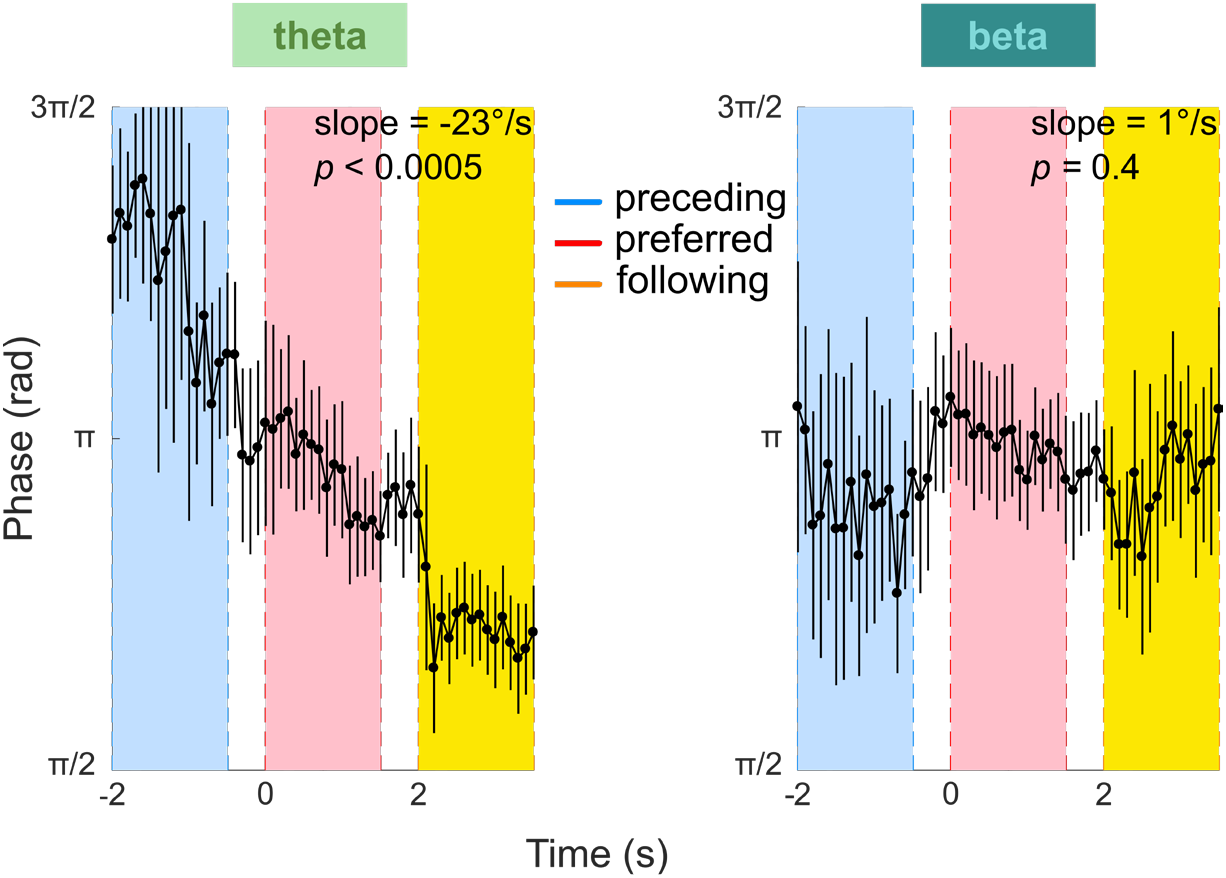
Phase transition over time in the theta (left) and beta (right) bands. The blue, pink and yellow shaded areas correspond to the stimulus periods of the preceding, preferred and following stimuli. The white regions between stimuli are inter-stimulus intervals, when the display rotates. The black dots and lines respectively correspond to the mean and standard deviation across cells. A negative phase slope was observed in the theta band (slope = -22.9 °/s, r^2^ = 0.92, *p* <0.00001), but not in the beta band (slope = 1.2 °/s, r^2^ = 0.02, *p* = 0.4).

The phase precession observed for the preceding, preferred and following stimuli was not observed for the remaining stimuli in the sequence, indicating that the phase selectivity did not extend to stimuli that far away from their preferred stimuli in the sequence. This decrease in phase locking is not unexpected if the preferred stimulus reappears every 5-7 stimuli, because items that follow the preferred stimulus will, at some point, be perceived to precede its next presentation, causing a breakdown of phase coding.

We carried out a number of control analyses to rule out (1) that the phase differences were caused by differences in firing rates or (2) differences in LFP power. First, we considered the possibility that a difference in the number of spikes fired for each stimulus was responsible for the phase differences (Figures S1 and S3). The largest spike-phase difference was observed between the preceding and following stimuli (p<0.0001), even though the firing rates were not significantly different from each other (Figure S1A; p=0.87). Furthermore, the preferred stimuli elicited a significantly higher firing rate than both preceding and following stimuli, yet their spike phase was intermediate. It is thus unlikely that a difference in firing rates could explain the phase precession effect. We also matched the spike count of individual trials between preceding and following stimuli, only including the subset of trials for the preceding and following stimuli in which the difference was <=1 spike (Figure S3) and observed that phase differences were maintained (F(1,1149) = 73.95; p<0.0001; Watson-William test, Bonferroni-corrected for multiple comparisons). We can therefore be confident that the differences in phase are a robust finding that is not caused by differences in the firing rate elicited by successive stimuli. Second, phase precession was also not caused by differences in LFP power, because LFP power was similar across the stimuli (Figure S1). Finally, we considered the possibility of a phase reset evoked by stimulus onset that could differentially affect spike-phase locking for the different stimuli (Figure S4). The absence of an influence of phase-reset was confirmed with a control analysis in which we removed precisely stimulus-locked spikes around the time of maximum phase reset (see Figure S4 and Methods).

Theta oscillatory activity in the primate ^17^ and human hippocampus ^18,19^ is often observed to be fragmented, with alternating periods of low and high theta power. We therefore asked whether the stimulus specific phase encoding reported above was influenced by variations in theta power. For instance, differences in oscillatory power have been suggested to modulate the “duty cycle” of phase coding ^20^. To address this question, we determined the median theta power for each recording channel. Spikes for each cell were then assigned to periods of low or high theta on the basis of the instantaneous theta power at the moment of the spike (median split). Within each period we then recomputed the phase of firing for the stimuli in the sequence. During both periods of low and high theta we found the same pattern of results as before: the neurons responded to the different stimuli at distinct theta phases, and the phase values shifted forward during the sequence (Figure S5). Thus, spike-theta phase coding was observed during periods of both high and low theta oscillatory activity.

## Discussion

Influential models of the hippocampus posit that this brain area structures incoming information by generating sequentially organized cell assemblies, each for a different input or event. In this scheme, a sequence of spatial locations (during navigation), or a sequence of items that has been learned, relies on the theta rhythm, which organizes the cell assemblies into sequences via phase precession ^10,21^. Our results provide important new insights into phase precession by showing (1) that phase precession occurs in the human MTL and (2) occurs outside navigation, in a task that required the learning of a sequence of visual objects. Neuronal firing in the human MTL elicited by different items in a sequence was phase-locked to the theta oscillation at distinct phases, and we observed a gradual phase precession that spanned three items in the sequence, with spikes fired at successively earlier phases as the sequence progressed. The phase coding was a robust phenomenon repeatedly observed in numerous control analyses: with a cross-validated selection of stimulus selectivity, when rejecting the spikes close (<0.6s) to stimulus onset, when equating the numbers of spikes between the preceding and following stimuli, and for both low and high theta power trials.

Recent studies in human and non-human primates have investigated how a series of items during memory tasks is encoded in different brain regions. At the level of single neurons, it appears that the last presented item is the most reliably encoded in firing activity ^2,3^, but these studies did not report phase locking to ongoing oscillations. Other studies have reported spike-phase coding with respect to higher frequency (32Hz) oscillations ^22^, or elevated theta-gamma phase-amplitude coupling ^23,24^. An MEG study Heusser et al., ^23^ is perhaps most relevant to the current study. They asked subjects to learn and recall the order of a sequence of images and showed that, on successfully encoded trials, consecutive items in the sequence exhibited higher gamma power at distinct theta phases. Our results go beyond by demonstrating theta-phase-coding of action potentials. This is important because phase-precession in rodents and, by extension, models of STM ^11^ and hippocampal function ^10^, posit a key role for neuronal spiking activity phase-locked to theta rhythms.

Theta oscillations and phase precession are prominent in the rodent hippocampus during spatial navigation ^9^. Spatial navigation in virtual environments in humans has revealed that navigational goals are represented in the firing activity ^25^. Theta oscillations have also been observed in humans, with frequencies ranging from 3-9 Hz ^8,13,14,26-28^. Watrous et al., (2018) ^8^ recently reported phase locking of action potentials to lower theta frequencies (3Hz) during navigation in a virtual environment. However, unlike our study, that study did not report a consistent directionality of theta phase coding with respect to the navigational trajectory of the subject, a difference that might be due to the different theta frequency range, or the different task. Here, we created a “conceptual” space, which consisted of a set of images that were ordered in a sequence and observed a distinct spike-phase code for successive items in the sequence.

The rate of phase precession in rodent place cells is not fixed, but varies as a function of the spatial extent of the place field, and the speed at which the animal crosses it ^29^. For example, when the same place field is crossed at slow or fast speeds (~12 cycles or ~5 cycles respectively), the phase shift from cycle to cycle slows down or speeds up accordingly. In our sequence, the number of stimuli that elicit elevated spiking is one to two stimuli ^4^. In other words, in our experiment, it takes ~24 cycles (2 seconds/stimulus for a 6Hz oscillation) to traverse the region in stimulus space associated with increased activity. This is a large number of cycles compared to the number that pass by when a rat traverses a hippocampal place field, and consequently the phase shift from cycle to cycle (or the phase slope reported in Figure 4) may be lower than that reported during spatial navigation in rodents. Future work could test how the rate of phase precession in a conceptual navigation space in humans varies as a function of the presentation speed of the sequence.

It is of interest to consider the response of multiple neurons (with distinct stimulus selectivities) within a single theta cycle. The finding that it is possible to track the phase advance of one neuron across theta cycles implies that a sequence of items is represented by a set of neurons tuned to different pictures within every theta cycle, firing at specific phases (Figure 5). During the cycle, neurons whose preferred stimulus is the previous item fire at the earliest phase and are followed by neurons coding for the current and following stimulus, analogously to the behavior of place cells.

**Figure 5:**
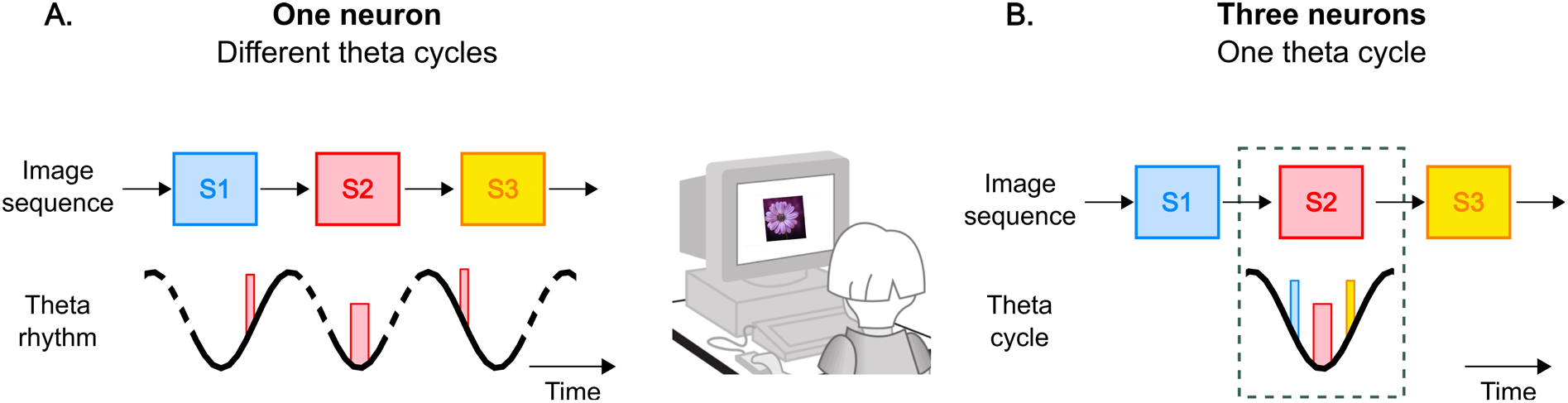
A subject views a sequence of three images (S1 to S3). A neuron whose preferred stimulus is S2 fires at the trough of the oscillation during the presentation of S2 (~180°; see Figure 3B). S1 and S3 are the preceding and following stimuli for this neuron and it fires at phases ~230° and ~120° when S1 and S3 are presented, respectively (Figure 3B). The thickness of the bars represents the firing rate. The stippled lines for the theta rhythm indicate that the theta cycles are not necessarily consecutive. B) While stimulus S2 is the preferred stimulus for one neuron, it is also the following stimulus for the neuron selective to S1, and the preceding stimulus for the neuron selective to S3. When S2 is presented on the screen, the neuron whose preferred stimulus is S2 fires around the trough of the oscillation. The neuron whose preferred stimulus is S1 (in blue) will fire at ~120° (S2 is the following stimulus for this neuron). Finally, the neuron whose preferred stimulus is S3 (in yellow) will fire at ~230° (S2 is the preceding stimulus for this neuron). Thus, in a given theta cycle, successive items are represented in the same order as the temporal order of the sequence.

In summary, learning and maintaining the order of a series of stimuli or events is crucial in many tasks. An accurate encoding of a sequence of ordered stimuli enables an organism to predict the future based on regularities learned in the past. Some authors have argued that the role of the hippocampus is to encode events that occur in a temporally organized sequence ^30,31^, by generating sequentially organized cell assemblies ^10,21^. Our results are broadly consistent with these hypotheses in that we observe distinct theta phase firing for successive items in a sequence, that reflects the sequence order. Nevertheless, the importance of phase coding is still under dispute, given that some species, such as bats, have excellent navigational capabilities and similar neuronal place coding strategies, which do not depend on the theta rhythm ^32-34^. The precise role of theta-phase precession therefore remains to be determined. It is encouraging that it is now possible to systematically study theta phase-coding in the human brain so that future studies can also use this approach to test the generality of theta-phase shifts during sequence coding, navigation in real and conceptual spaces, STM, and other cognitive functions.

## Methods

Participants were nine patients (four females, age range 18-36 years) with pharmacologically intractable epilepsy undergoing epileptological evaluation at the Amsterdam University Medical Center, location VUmc, The Netherlands. Patients were implanted with chronic depth electrodes for 7-10 days in order to localize the seizure focus for possible surgical resection ^35,36^. All surgeries were performed by J.C.B and S.I. The Medical Ethics Committee at the VU Medical Center approved the studies. The electrode locations were based entirely on clinical criteria and were evaluated based on the pre-surgical planned trajectories on the basis of structural MRI scans. The accuracy of the implantation was always checked using a CT scan co-registered to the MRI. We only included electrodes that were within a 3mm deviation from the target (based on visual confirmation).

Each electrode contained eight microwires (Behnke-Fried electrodes, Ad-Tech Medical) from which we recorded single/multi-unit activity and local field potentials, and a ninth microwire that served as a local reference. The signal from the microwires was recorded using a 64-channel Neuralynx system, filtered between 1 and 9000 Hz, sampled at 32KHz. On average, each patient was implanted with 34 ± 11.8 microwires. Participants sat in their hospital room at the Epilepsy Monitoring Unit, and performed the experimental sessions on a laptop computer. All patients participated in the two types of experimental sessions described below.

### Sequence Learning (SL) Sessions

The patients performed a total of 27 sequence learning (SL) sessions. In each SL session, subjects were presented with a sequence of 5-7 images (image number determined as a function of the difficulty level and the availability of the patient). The images were always in a pre-determined order such that a given image, A, predicted the identity of the next image, B, and so on. Subjects were asked to remember the order of the stimuli in the sequence. Each stimulus was presented for 1.5s with an inter-stimulus interval (ISI) of 500ms, resulting in individual trials of 2000ms (Figure 1). The sequence was repeated continuously 60 times resulting in experimental sessions of ~10-14 minutes, not including time spent by the subject to respond on probe trials. 20% of trials were “probe” trials in which, instead of being presented with the next image of the sequence, subjects were shown two images side by side and asked to decide (by pressing one of two keys on the keyboard) which of the two was the next image in the sequence.

To further the impression of a sequence of images we used the following display arrangement (Figure 1): Each image was presented at the center of the screen while placeholders (empty gray squares) were presented to the left and right of the central image. At the end of the 1500ms presentation period, the central image was replaced by a gray placeholder and all the gray squares moved one “step” forward in a clockwise direction for the duration of the ISI, such that each placeholder eventually occupied the next placeholder position. At the end of the ISI, the placeholder that now occupied the central position was replaced by the next image in the sequence. The viewer’s subjective impression at the end of the ISI interval was that the central image had been hidden, and then moved clockwise, while the central position was replaced by the next image in the sequence.

### Spike Detection and Sorting

Spike detection and sorting were performed with wave_clus ^37^. Briefly, the data were band pass filtered between 300-3000Hz and spikes were detected with an automatic amplitude threshold (Figure S6). Spike sorting was performed with a wavelet transform that extracted the relevant features of the spike waveform. Clustering was performed using a super-paramagnetic clustering algorithm. Clusters were visually reviewed by the first-author for 1) the mean spike shape and its variance; 2) the ratio between the spike peak value and the noise level; 3) the inter spike interval distribution of each cluster; 4) the presence of a refractory period for the single-units; i.e. fewer than 1% of spikes in a 3ms or smaller inter-spike interval; 5) the similarity of each cluster to other clusters from the same microwire. Based on manual inspection of these criteria, clusters were retained, merged or discarded. Unit quality metrics are shown in Figure S7.

### Number of neurons and their locations

Over the nine patients we recorded from 619 neurons (single and multi-unit) in the left and right hippocampi (*N*=491), temporal cortices (*N*=110), and parahippocampal cortices (*N*=18) in the sequence learning sessions. In the main analyses, all cells were pooled together. We did not notice differences in theta power between the hippocampal and parahippocampal/temporal electrodes (two-sample t-test, p = 0.56; mean ± SEM = 21 ± 1 vs. 22 ± 1 uV), or in the strength of phase locking as measured by the kappa concentration parameter ^38^ (F =1.0, p=0.98). Additionally, when the main analyses were performed on only hippocampal cells, the same pattern of stimulus specific phase-locking was observed as in the main analysis (all p-values < 0.0001; Watson-William test for equal means, Bonferroni-corrected for multiple comparisons).

### Data Analysis

All analyses were performed with the FieldTrip toolbox ^39^ and custom Matlab code.

## Estimation of theta-band oscillatory activity

All LFP analyses were performed with the FieldTrip toolbox in Matlab ^39^. The LFP was recorded from the same microwires as the spiking activity. It was downsampled to a 1,000-Hz sampling rate and notch filtered (between 45-55Hz and 98-102Hz) using a second order Butterworth filter. For each channel, we computed the time-frequency decomposition for 28 different frequencies: f = 2^x^ with x ∈ {6/8, 8/8, 10/8,…60/8} ^7^. The time frequency decomposition was performed with the multitaper method over a 3 second epoch encompassing the preceding and preferred stimuli of the raw LFP trace, using two cycles per time-window at each frequency. We estimated whether significant theta activity was present in the LFP by fitting a 1/f function to the power spectrum and taking the ratio (in units of decibels) between the actual power spectrum and the 1/f fit (Figure 2A). Significance was estimated with a t-test, Bonferroni corrected for multiple comparisons (28 frequencies). In addition to estimating theta power with the raw traces, we also measured the power spectrum of the oscillations around the time of each spike (spike triggered power; Figure 2B). We extracted a 1s LFP segment centered on each spike and extracted the frequency spectrum of each segment. The average power spectrum of these LFP traces was estimated by taking the average of the absolute values (the power) of the spectra of all LFP segments ^40^. The resultant power spectrum was fitted to a 1/f function, and the ratio (in decibel units) computed. Significance was estimated with a t-test, Bonferroni corrected for multiple comparisons over 28 frequencies. Both measures of quantifying oscillatory power revealed prominent theta activity in the 4-8Hz range and beta activity in the 10-18Hz range.

## Estimation of spike-LFP phase-locking

The LFP phase was estimated using a Hanning taper over the entire duration of the session (i.e., without epoching), using five cycles per time-window at each frequency. A phase of zero corresponds to the peak of the LFP oscillation, and a phase of ± 180 corresponds to the spike being at the trough of the oscillation. Phase-locking was evaluated by comparing the distribution of phase angles against the uniform distribution using the Rayleigh test. We observed significant phase locking in the 4-8 Hz (theta range) and 10-18 Hz (beta range). For subsequent analyses, the LFPs were therefore band-pass filtered with a second order Butterworth filter in these frequency ranges. A phase value for each spike in each frequency range was extracted using the Hilbert transform on the band-passed signal.

## Stimulus specific spike-phase-locking

To determine whether the spikes for the different stimuli occurred at different phases, we assigned phase values to each stimulus depending on the time at which the spikes were fired. Phase values were binned into 15 bins. For visualization purposes only, a smoothing of two bins in both directions was applied. To compare the phase distributions between stimuli the Williams-Watson test from the Circular Toolbox for Matlab ^38^ was applied to the unsmoothed data.

The preferred image of a neuron was the stimulus that elicited the most spikes. The images before and after the preferred image in the sequence were designated as the preceding and following images, respectively. The phase analysis was also performed using k-fold cross-validation. For each neuron, we determined the preferred, preceding, and following stimuli on all trials except one. The phase values for these stimuli were then extracted on the left-out trial and all the analyses were performed on the data of the left-out trial.

To control for the effect of spike number on phase between the following and preceding stimuli (Figure S3), we equalized the number of spikes for each neuron by only including the subset of trials in which the difference on each trial between the two stimuli was <=1 spike. We re-computed the preferred phases for the preceding and following stimuli, while only considering the reduced number of trials.

## Phase-Reset and Inter-trial Coherence (ITC)

We controlled for the possible influence of phase-reset of the theta oscillation caused by stimulus onset on measures of phase precession (Figure S4). We computed the inter-trial coherence at each time and frequency point. On each trial of the following, preferred and preceding stimuli we extracted an LFP segment in the time window of [-0.5 1.5] sec (time 0 corresponds to stimulus onset). The phase and power at each time and frequency point was extracted from a time frequency transform of the signal. The parameters for the time-frequency transform are the same as described above: namely, the multi-taper method with 2 cycles per frequency on the notch-filtered and down sampled signal, at 28 frequencies, and in a time interval of -0.5s to 1.5s in steps of 50ms. The inter-trial coherence is the absolute value of the average spectrum normalized by its amplitude ^41^, and varies between zero (no phase-locking) and one (perfect phase-locking). Equal numbers of trials were used for the different stimuli.

### Statistical Testing

We used two-tailed tests unless otherwise specificed. All statistical tests performed on circular data were performed with the Circular Toolbox for Matlab ^38^, applied to the unsmoothed data.

### Data Availability

The data supporting the findings of this study are available from the corresponding author upon reasonable request.

### Code Availability

The code supporting the findings of this study are available from the corresponding author upon reasonable request.

### Author Contribution

L.R and P.R.R designed the study. J.C.B and S.I performed the surgeries. M.S, B.Z, M.P, J.K.P, J.C.P and L.R. collected data. L.R analyzed the data with input from R.V. L.R. wrote the first version of the manuscript. L.R and P.R.R finalized the manuscript. All authors commented on the finalized version of the manuscript.

## Acknowledgments

We are grateful to to C.J. Stam, E. van Dellen, L. Douw, P. Ris, S. Claus, D. Velis, and R. Joosten for help with obtaining ethics approval and for technical help with the patients and recordings. This work was supported by grants from the French Agence Nationale de la Recherche (ANR-12-JSH2-0004-01), the Fyssen foundation, and the Université Paul Sabatier, Toulouse, France (BQR, 2009 and Appel à Projets de Recherche Labellisés, 2013), to L.R., the European Research Council (ERC Consolidator Grant P-Cycles number 614244) and an ANITI (Artificial and Natural Intelligence Toulouse Institute) Research Chair (ANR-19-PI3A-0004) to R.V., the Studienstiftung des Deutschen Volkes (German Academic Scholarship Foundation) to B.Z., the European Union (ERC Grant Agreement n. 339490 “Cortic_al_gorithms” and grant agreements 720270 and 785907 ‘‘Human Brain Project SGA1 and SGA2’) and the Friends Foundation of the Netherlands Institute for Neuroscience to P.R.R.

