## Supplemental Material for "Theta-phase dependent neuronal coding during sequence learning in human single neurons"

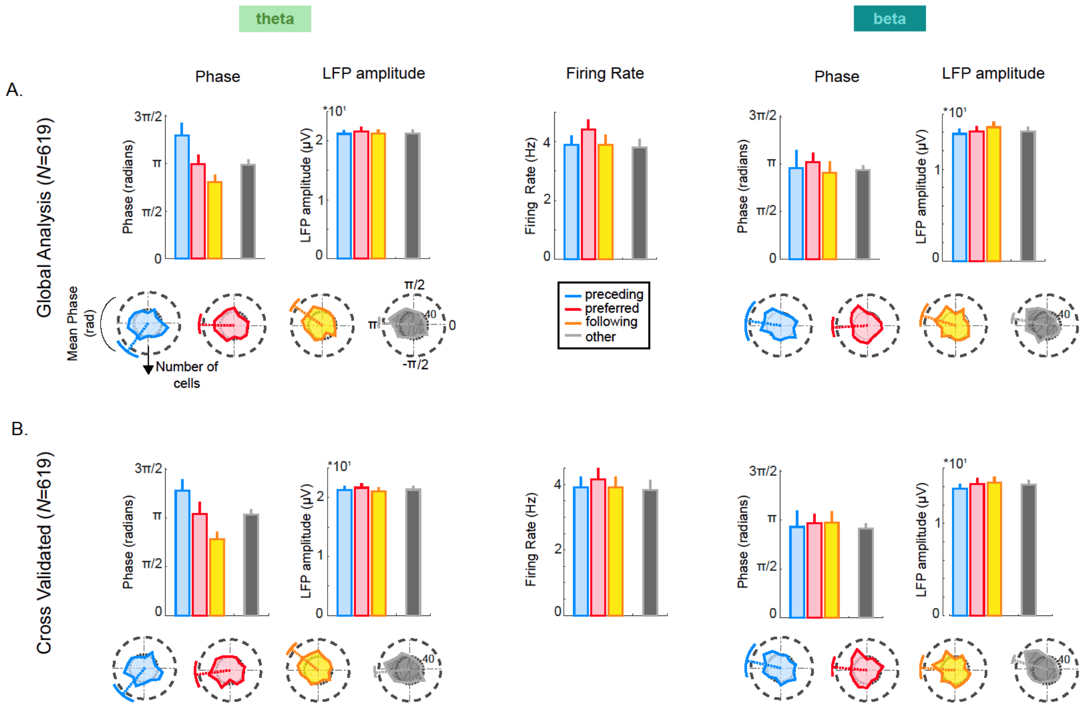


Figure S1. Differences in firing rates and LFP amplitudes do not account for phase precession. Phase, LFP amplitude (square root of LFP power), and firing rates are shown for the preceding (blue), preferred (red), following (yellow), and other (black) stimuli, in the theta (left column) and beta (right column) bands. For each neuron a preferred stimulus was selected and the stimuli before and after the preferred stimulus in the sequence were labeled as the preceding and following stimuli. Since the number of stimuli in the sequence varied across subjects (between 5-7; as a function of the difficulty level and availability of each patient), the number of other stimuli in the sequence also varied. These remaining stimuli in the sequence are therefore grouped together as the "other" stimuli in this figure (black). The bar graphs showing phase values show the same data as the rose plots (in a different format). In all panels, error bars correspond to the *SEM* across cells.

A) Global analysis. For each neuron (*N* = 619) the preferred image was the one that elicited the most spikes during the experiment. The largest phase difference was observed for the preceding and following stimuli in the theta band even though their firing rates and LFP amplitudes were not different (phase difference, Watson-William Test, $F(1,1237) = 102.05$; $p<0.0001$, Bonferroni corrected for multiple comparisons; firing rate difference, $p=0.9$; LFP amplitude difference, $p=0.8$). The blue, red and yellow rose plots are the same as in Figure 3B, D. The firing rate for the "other" stimuli was significantly lower than for the preceding and following stimuli ($p<0.005$). The phase difference between the "other" stimuli and the preceding and following stimuli was significant ($p<0.0001$) in the theta band, but not in the beta band.

B) Cross-validated analysis. For each neuron (*N* = 619) the preferred, preceding, following, and "other" stimuli were chosen on the basis of all trials but one. The phase values, firing rates, and LFP amplitudes for these stimuli were obtained on the left-out trial. The procedure was repeated for each trial of the experiment. In the cross-validated data, the same pattern of phase differences, and the order of phases, as in Figure 3 were observed. The phase precession was significant ($F(1,1241) = 24.02$; $p<0.0001$ for preferred vs. preceding; $F(1,1241) = 29.4$; $p<0.0001$ for preferred vs. following; and $F(1,1241) = 105.18$; $p<0.0001$ for preceding vs. following; $F(1,1241) = 27.28$; $p < 0.0001$ for other vs. preceding; $F(1,1241) = 34.81$; $p < 0.0001$ for other vs. following; Watson-William test Bonferroni corrected for multiple comparisons). The phase differences were not significant in the beta band. The firing rates for the preceding and following images were not significantly different from each other ($p=0.96$), but the firing rate differences between all other stimulus pairs were significant ($p<0.05$). Finally, the theta and beta LFP amplitudes were not significantly different between stimuli ($F(3,2483)=0.13$; $p = 0.94$ for theta, $F(3,2483)=0.22$; $p = 0.88$ for beta).

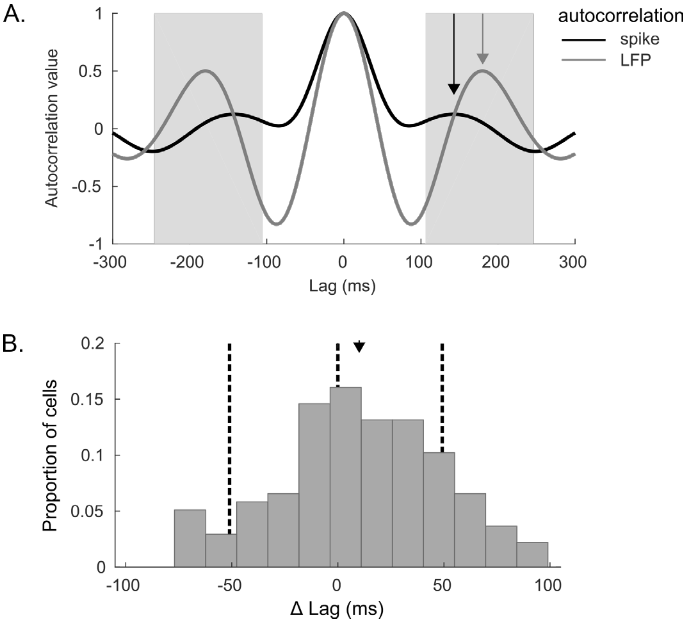


Figure S2. A. Autocorrelations of spikes and LFPs. A) Representative example of the autocorrelation of a neuron and the LFP from the same recording site. The autocorrelation function was computed for the spikes (smoothed with a 25ms standard deviation Gaussian window) and for the LFP filtered in the theta-band. Peaks in the autocorrelations were detected in a time window of ± 70ms around the peak of the average LFP auto-correlogram across all cells, which was 178ms (this window has been indicated in grey). Note that for this cell, the first side peak for the spikes (black arrow) occurs at a shorter lag than the first side peak of the LFP autocorrelation function. In other words, the frequency of the oscillation is slightly higher for the spikes, as is expected during phase precession. B) Distribution of the difference between the peak lag of the LFP and the peak lag of the spikes across cells. The mean of the distribution (indicated by the black arrowhead) is shifted significantly to the right of 0 (mean ± sem = 10.07 ± 3.3 ms; 95% confidence interval of the mean = [3.6 16.6], $t(1,136) = 3.05$, $p < 0.005$). To increase the reliability of the measure, only cells (*N*=137) that showed a clear theta-band peak in the autocorrelograms for both spikes and LFPs were included (i.e., the difference between the detected peak and the minimum of the autocorrelation in the highlighted time window was greater than a threshold of 0.15).

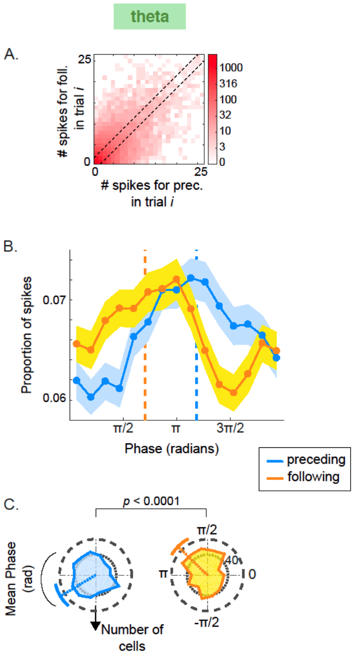


Figure S3. Control for the possible influence of firing rate differences on phase precession measures. A. The spike count elicited by preceding and following stimuli was matched at the level of individual trials. Specifically, we only included trials in which the firing rate difference was <=1 spike. The density plot shows the number of spikes fired in response to the preceding and following stimuli on each trial *i*. The main analyses (Figures 3, 4 and S1) used all trials. The analysis presented in this figure only uses trials that lie between the two dashed diagonal lines, i.e., trials on which the difference between the two stimuli was <=1 spike (N=575 cells remained). The mean and standard deviation of trials per cell was 13 ± 7 Hz (as opposed to 24 ± 5 Hz for the main analysis; Figure 3).
B. Distribution of spike phases relative to the theta band LFP for the preceding (blue), and following (yellow) stimuli when using trials with matched firing rates (between the two dashed diagonal lines in panel A; $N = 575$).
C. Firing phase across all cells *(N = 575),* with colors as in panel (B). The format of panels B and C is identical to that of Figure 3 (now only showing preceding and following stimuli, for which the spike numbers have been equated).

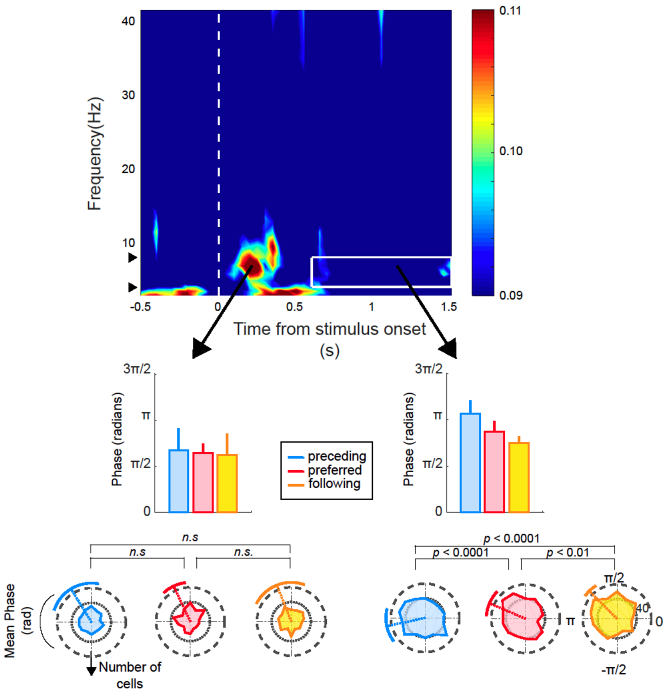


Figure S4**.** Influence of phase-reset and inter-trial coherence (ITC) on phase precession. (*top*) The ITC quantifies the degree of phase locking of the LFP to stimulus onset. It was computed at each time and frequency point with respect to the onset of the preceding, preferred, and following stimuli. The black arrowheads on the y-axis correspond to the 4-8Hz frequency band. The phase reset caused by stimulus onset is visible as an increase in ITC, with a peak around 7Hz, at 250ms.

(*bottom left)* The phase calculated in a 60ms window around peak ITC is not significantly different between the preceding, preferred and following stimuli ($p > 0.1$).

*(bottom right)* Phase precession in a time window after 600 ms post-stimulus onset, during which ITC had subsided. In this window, phase precession was significant ($F(1,1225) = 16.1$; $p<0.0001$ for preferred vs. preceding; $F(1,1225) = 6.8$; $p<0.01$ for preferred vs. following; and $F(1,1225) = 41.4$; $p<0.0001$ for preceding vs. following; Watson-William test).

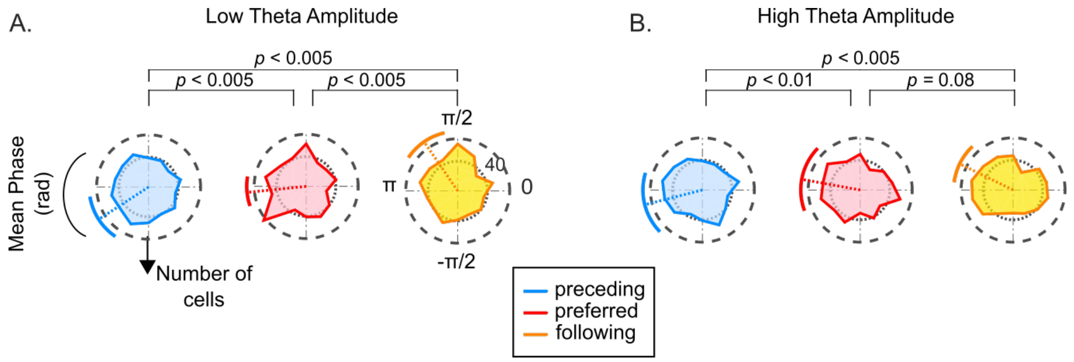


Figure S5. Spike-phase coupling during periods of A) low and B) high theta amplitude. For each cell, each spike was labeled as occurring during periods of either low or high theta relative to the median of the distribution of theta amplitudes of the recording channel across trials. The rose plots show the distributions of spike phases for the preceding (blue), preferred (red) and following (yellow). Phase precession was evident during epochs of low and high theta (compare to Figure 3). Low theta: $F(1,1237) = 9.9$; $p<0.005$ for preferred vs. preceding; $F(1,1237) = 51.5$; $p<0.0001$ for preferred vs. following; and $F(1,1237) = 98.32$; $p<0.0001$ for preceding vs. following; Watson-William test. High-theta: $F(1,1237) = 7.18$; $p<0.01$ for preferred vs. preceding; $F(1,1237) = 2.98$; $p=0.08$ for preferred vs. following; and $F(1,1237) = 19.96$; $p<0.0001$ for preceding vs. following.

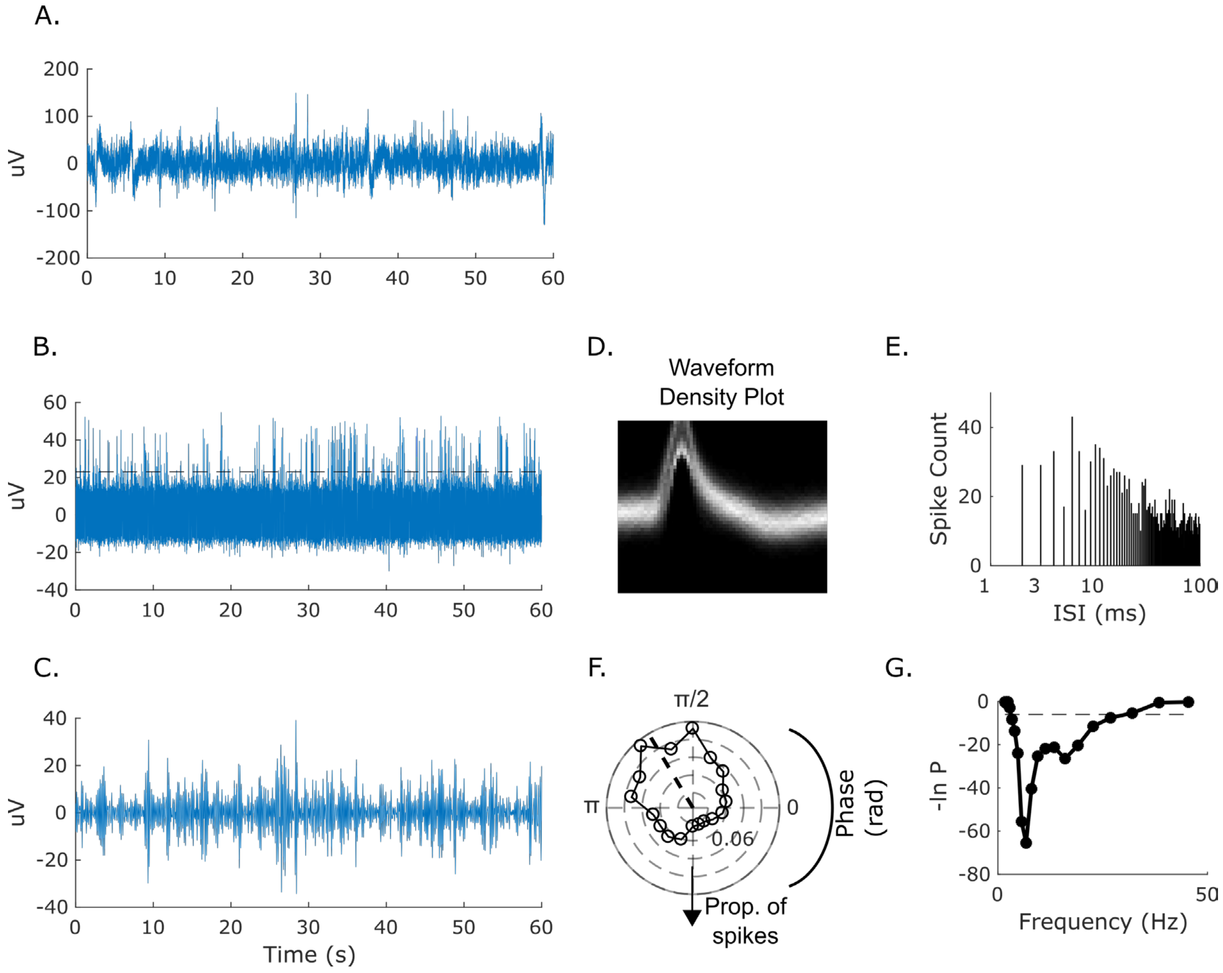


Figure S6. Signal recorded from an example microwire. A) Unprocessed signal recorded on one microwire in an epoch of 60s. B) The signal was filtered between 300-3000Hz to extract spikes. The dashed black line corresponds to the threshold used for spike detection. C) The LFP was bandpass filtered between 4-8Hz to extract the phase of each spike. D) Density plot of the spike waveforms. E) Inter-spike interval distribution. F) Phase distribution of spikes in the 4-8Hz theta band. The dashed black line indicates the mean phase for this cell. G) Significance of phase locking (Rayleigh test). The threshold for significance (horizontal black dashed line) is set to p<0.05, Bonferroni-corrected for multiple comparisons. The shaded gray area corresponds to the theta band (4-8Hz). It can be seen that this cell showed significant theta phase locking.

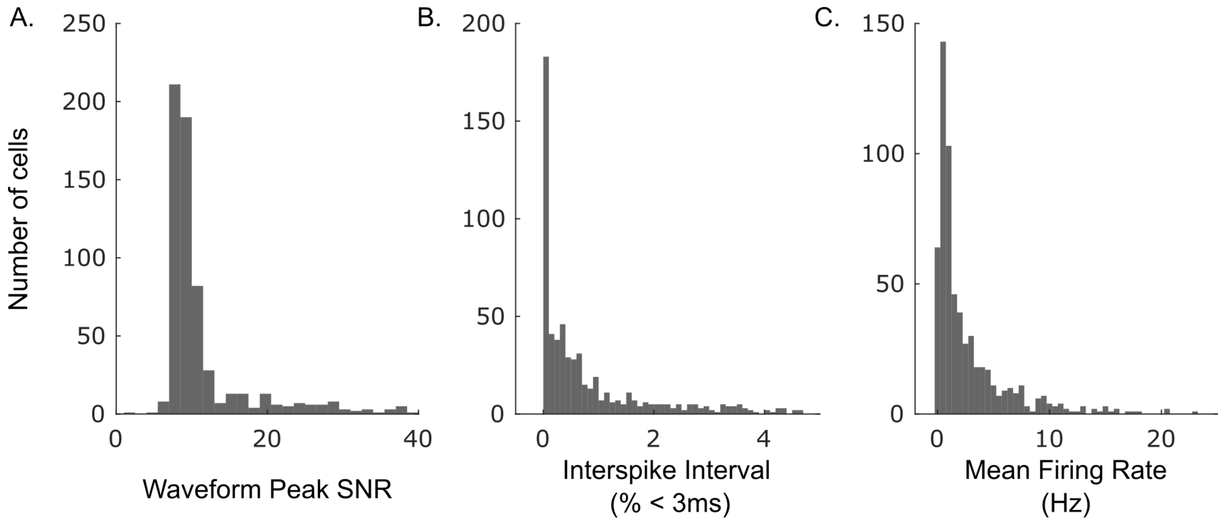


Figure S7. Spike quality metrics. A). Histogram of the SNR of the peak of the mean waveform of each unit. B). Histogram of the percentage of interspike intervals that are < 3ms. Note that the large majority of units had less than 1% of interspike intervals < 3 ms. C). Histogram of the mean firing rates.
